# Functional comparison of full-length palladin to isolated actin binding domain

**DOI:** 10.1101/2022.09.14.507982

**Authors:** Sharifah Albraiki, Rachel Klausmeyer, Oluwatosin Ajiboye, Moriah R. Beck

**Affiliations:** Department of Chemistry and Biochemistry, Wichita State University, Wichita, KS 67260, United States; Department of Chemistry and Geosciences, Jacksonville State University, Jacksonville, AL 36265, United States

**Keywords:** G- and F-actin binding, polymerization, palladin, autoinhibition

## Abstract

Palladin is an actin binding protein that is specifically upregulated in metastatic cancer cells but also co-localizes with actin stress fibers in normal cells and is critical for embryonic development as well as wound healing. Of nine isoforms present in humans, only the 90 kDa isoform of palladin, comprising three immunoglobulin (Ig) domains and one proline-rich region, is ubiquitously expressed. Previous work has also established that the Ig3 domain of palladin is the minimal binding site for F-actin. In this work, we compare functions of the 90 kDa isoform of palladin to the isolated actin binding domain. To understand the mechanism of action for how palladin can influence actin assembly, we monitored F-actin binding and bundling as well as actin polymerization, depolymerization, and copolymerization. We also provide initial evidence that 90 kDa palladin exists in a closed conformation that prevents binding by the Ig3 domain to G-actin as compared to the isolated domain. Understanding the role of palladin in regulating the actin cytoskeleton may help us develop means to prevent cancer cells from reaching the metastatic stage of cancer progression.

## 1 INTRODUCTION

Palladin localizes with actin-rich structures in normal cells, but increased expression is also correlated with the metastatic stage of cancer cells (Welsch et al. 2007; Goicoechea et al. 2009). Palladin is an actin crosslinking protein that also acts as an actin scaffolding molecule by interacting with a large number of actin regulatory proteins (Dixon et al. 2008; Vattepu et al. 2015). Nine different palladin isoforms have been identified, where the largest isoform (iso1 or 200 kDa) contains five immunoglobulin domains and two proline rich regions. Only two isoforms (iso3 and iso4) have been reported to be expressed in myofibroblasts(Ronty et al. 2006); however, the role of these isoforms remains unclear. In this study, we compared the roles of the isolated actin binding domain of palladin (Ig3) with that of the 90 kDa or isoform 4 which is the only ubiquitously expressed isoform that is also upregulated in cancer cells. The 90 kDa palladin (90 kDa-Palld) isoform comprises the three C-terminal immunoglobulin domains (Ig3, 4, and 5) and one proline rich region (PR2). This second proline rich region binds to many actin-regulatory proteins such as VASP (Brindle et al. 1996; Boukhelifa et al. 2004), profilin (Boukhelifa et al. 2006), Eps8 (Goicoechea et al. 2006), α-actinin (Ronty et al. 2004), Lasp-1 (Rachlin et al. 2006) and Ezrin (Khanna et al. 2004). Palladin also binds directly to other proteins that indirectly influence actin organization such as ArgBP2, LPP and SPIN-90 (Ronty et al. 2005; Ronty et al. 2007). Recent investigation of 90 kDa-Palld suggested that this isoform in particular also plays a role in promoting the inflammatory function of cancer-associated fibroblasts (CAFs) highlighting the therapeutic potential of targeting CAFs to prevent cancer metastasis (Alexander et al. 2021).

The Ig3 domain of palladin is the minimal binding site of actin and has been the focus of much of the previous biochemical characterization of palladin (Dixon et al. 2008). Using this isolated Ig3 domain of palladin (Ig3-Palld), the Beck Lab has established that it promotes actin assembly and lowers the disassembly rate of actin filaments (Gurung et al. 2016). Ig3-Palld has also been shown to dimerize upon interaction with F-actin which supports the bundling and cross-linking of actin filaments (Vattepu et al. 2015). Interestingly, although the Ig4 domain of palladin does not bind directly to F-actin, the tandem Ig34 domain binds more tightly than Ig3 alone (Beck et al. 2013). This is echoed by a study that established that the membrane phosphoinositide phosphatidylinositol (PI) 4,5-bisphosphate [PI(4,5)P_2_] binds to palladin through the Ig3 domain thereby influencing palladin behavior and reducing its ability to polymerize and crosslink actin. The isolated Ig4 domain shows only minimal binding to PI(4,5)P_2_, but the Ig34 tandem domain binds more tightly than Ig3 (Yadav et al. 2016). The mechanism behind this affect has yet to be explained, but it could be due to interactions between the Ig3 and Ig4 domains or an unknown function of the unique, extended linker region that connects the two domains.

Previous data has also shown that Ig3-Palld and 90 kDa-Palld play different roles in organizing the actin cytoskeleton *in vivo*. Overexpression of 90 kDa-Palld *in vivo* can replace the Arp2/3 complex in nucleating actin filaments in the presence of the Arp2/3 inhibitor CK-666 (Dhanda et al. 2018). Palladin also has the ability to maintain the actin-based motility of *Listeria monocytogenes* in the absence of the Arp2/3 complex. Biomimetic assays revealed that 90 kDa-Palld was able to nucleate actin clouds around the bacteria and enhanced the formation of actin-based comet tails *in vitro*; however, the formation of the actin cloud and comet tails did not occur when the Arp2/3 complex was substituted with Ig3-Palld (Dhanda et al. 2018). While the mechanism of action for how palladin contributes during the bacterial invasion is not fully understood, previous data has revealed that Ig3-Palld binds to F-actin via several surface exposed lysine residues (Beck et al. 2013). Mutations of lysine residues (K15A, K18A, and K51A) in the Ig3 domain nearly abolish all F-actin binding *in vitro* and disrupt cellular actin when expressed *in vivo*. Furthermore, mutation of these same lysine residues in 90 kDa-Palld results in tortuous, shortened actin comet tails and a significant decrease in *L. monocytogenes* speed (Dhanda et al. 2018). This finding indicates that these lysine residue mutations in Ig3-Palld diminish the binding to actin and prevent palladin from crosslinking actin filaments which results in disoriented and warped comet tails.

In this work we compare the different activities and functions of Ig3-Palld and 90 kDa-Palld directly. While 90 kDa-Palld has never previously been purified from bacterial cells due to instability and low yield, here we have successfully purified 90 kDa-Palld from *E. coli* and now can test the effect of this protein on actin structures and functions using biochemical assays. Our data indicates that 90 kDa-Palld binds tightly and crosslinks F-actin but does not interact significantly with G-actin as opposed to the isolated Ig3-Palld domain. Our data also suggest that 90 kDa-Palld may be autoinhibited thereby blocking the G-actin binding site and preventing the binding of 90 kDa-Palld to monomeric actin.

## 2 RESULTS

### 2.1 Purification of the 90 kDa isoform of palladin

After many previous unsuccessful attempts to purify 90 kDa-Palld from *E. coli*, the human gene was cloned into a vector containing a His_6_-tag and TEV cleavage site on the N-terminus and successfully expressed in *E. coli* in high yield. Several modifications were implemented to the Ig3-Palld protein purification protocol to improve the expression and purification of 90 kDa-Palld. In particular, autoinduction media was used for expression instead of induction with IPTG in LB media (Studier 2005). Additionally, all purification steps were performed at 4 °C and 10% glycerol was added to all purification buffers to stabilize the 90 kDa-Palld. (Supplementary Figure S1)

### 2.2 Comparison of actin binding and crosslinking

The ability to successfully purify 90 kDa-Palld from bacterial cells provides the opportunity to investigate the mechanism of action of full-length palladin on actin regulation *in vitro*. In this study we first determined whether the 90 kDa-Palld purified from bacterial cells behaves similarly to the one isolated from insect cells (Table 1). In addition, we compared the effect of the 90 kDa-Palld with the isolated Ig3-Palld domain on actin organization and dynamics. To compare the actin-binding affinity of the 90 kDa-Palld with that of Ig3-Palld, we carried out actin co-sedimentation assays where the concentration of palladin was held constant at 10 μM and increasing amounts of F-actin were used. Representative gels and the average binding curve and associated linear fits for the assay are shown in Figure 1. The bacterially expressed 90 kDa-Palld was shown to have nearly identical binding affinity to 90 kDa-Palld expressed in insect cells (Dixon et al. 2008). As shown in Table 1, the apparent dissociation constant obtained in this study for 90 kDa-Palld (K_d_=2.11 +/-1.09 µM) was slightly lower than the isolated Ig3-Palld domain (K_d_= 3.42 +/-1.24 µM) indicating that 90 kDa-Palld has a greater affinity for F-actin than Ig3-Palld. The approximate K_d_ for Ig3-Palld reported by Dixon *et al*. was hampered by a limitation in the Ig3 concentration range necessary for acquiring a saturation curve (Dixon et al. 2008). In our experiments, we circumvented this issue by varying the concentration of F-actin instead of palladin and were able to obtain a good saturation curve, which may explain why our binding affinity was quite different than that previously reported.

**TABLE 1.** COMPARISON OF EQUILIBRIUM DISSOCIATION CONSTANTS Apparent equilibrium dissociation constants, K_d_, of Ig3-Palld and 90 kDa-Palld from bacterial (*E. coli*) cells and insect (Sf9) cells.

| Protein | K<sub>d</sub> (μM) | B<sub>max</sub> |
| --- | --- | --- |
| **Ig3-Palld(Dixon et al. 2008)\*** | ~ 60 | N.D. |
| **Ig3-Palld (this work)** | 3.42 ± 1.24 | 3.95 |
| **90 kDa-Palld (*E. coli*)** | 2.11 ± 1.09 | 6.27 |
| **90 kDa-Palld (Sf9)(Dixon et al. 2008)** | 2.1 ± 0.5 | N.D. |

\*Binding data for previously reported Ig3 K<sub>d</sub> was an estimate limited by lack of complete saturation curve.

**Figure 1.**
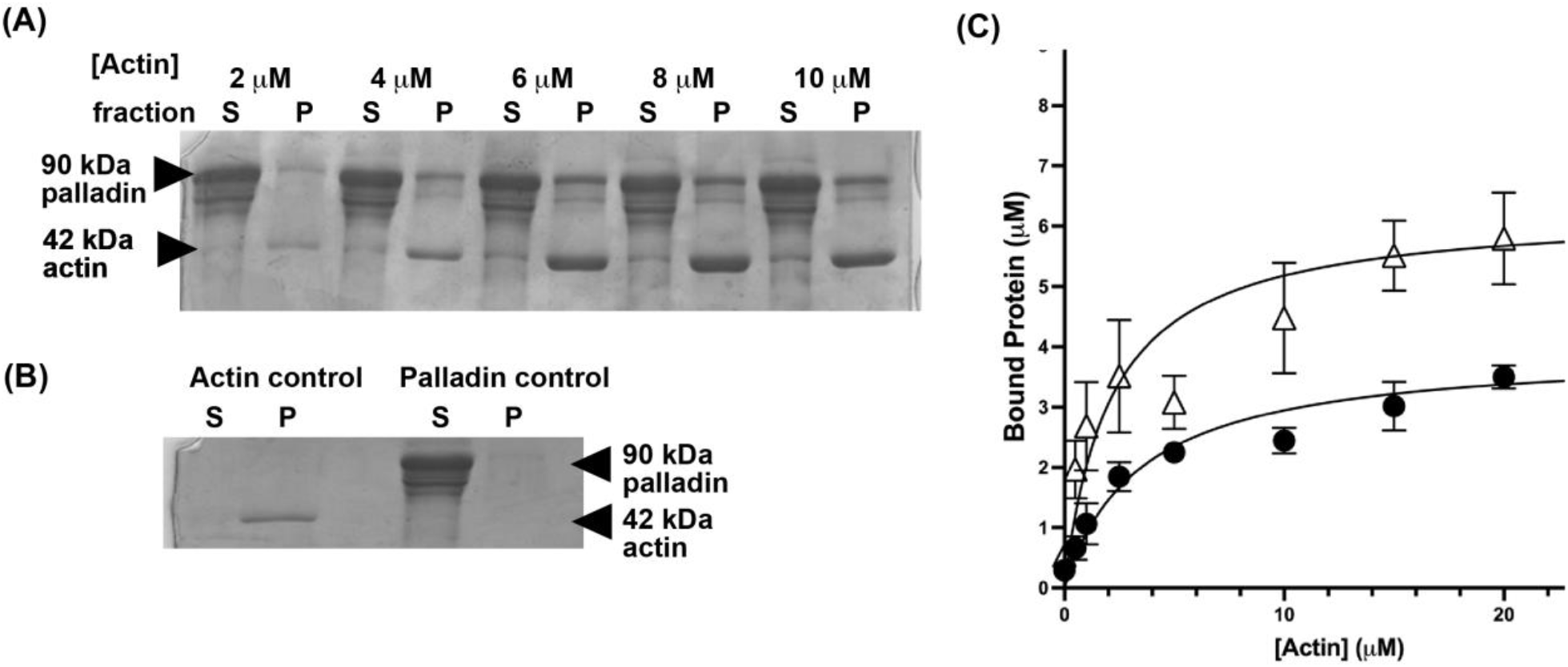
F-actin co-sedimentation assay to compare 90 kDa-Palld and Ig3-Palld binding to F-actin. (A) Representative SDS-PAGE gel of co-sedimentation assay with supernatant fractions (S) and pellet fractions (P) of a constant concentration of 90 kDa-Palld (10 µM) and various concentrations of actin (0-10 µM). (B) Also shown are actin and palladin controls. (C) The fraction of palladin that co-sediments with F-actin was monitored by densitometry of the SDS-PAGE gel. A hyperbolic curve was used to fit the data, assuming a 1:1 stoichiometry and specific binding only, which gives a dissociation constant for full-length palladin with F-actin of 2.11 ± 1.09 µM (triangles) and for the Ig3 domain of 3.42 ± 1.24 µM (closed circles). Data are means +/- standard deviation for three or more separate measurements.

To determine the bundling or crosslinking efficiency of 90 kDa-Palld, we once again turned to actin co-sedimentation assays where in this case the concentration of actin was held constant at 10 μM and increasing amounts of palladin were used (Figure 2). Based on a comparison of the data with that previously published for the isolated Ig3 domain of palladin (Dixon et al. 2008), 90 kDa-Palld has a significantly higher bundling efficiency than Ig3-Palld at lower concentrations (5 and 10 µM), but multiple unpaired t-test analysis of bundling at 20 µM palladin indicate that bundling is not significantly different at this concentration.

**Figure 2.**
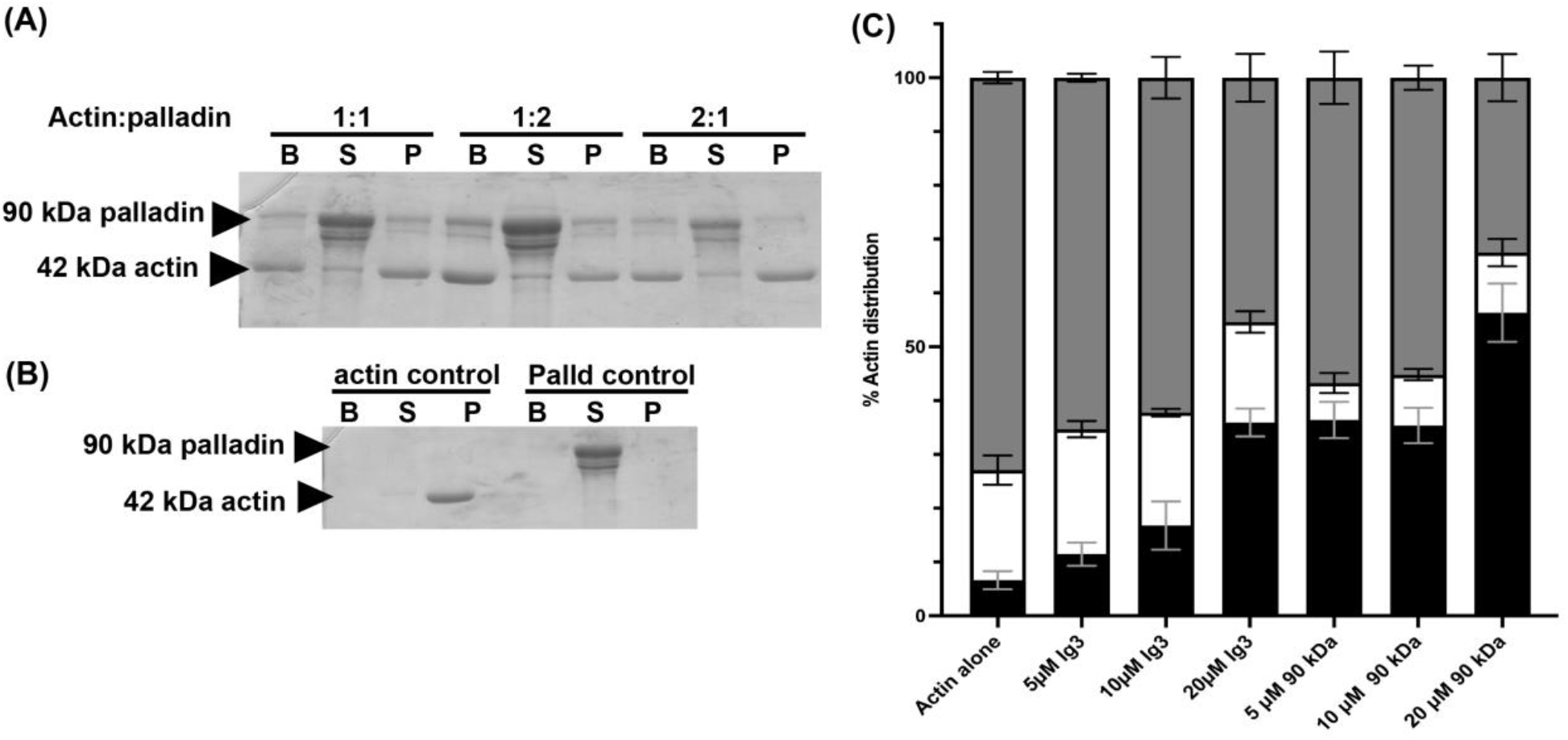
F-actin crosslinking by 90 kDa-Palld. (A) Representative SDS-PAGE gel of co-sedimentation assay with bundle (B), supernatant (S) and pellet fractions (P) of a constant concentration of actin (10 µM) and various ratios of actin to 90 kDa-Palld (1:1, 1:2, and 2:1). (B) An SDS-PAGE gel of actin and palladin controls (C) The percentage of actin found in the low-speed bundle (black), supernatant monomers (white), and pellet filaments (gray) recovered from incubation with either Ig3 or 90 kDa palladin was quantified with ImageJ. Data are means +/- standard deviation for three or more separate measurements.

### 2.3 Co-polymerization of actin and palladin

In the previous co-sedimentation assay, actin was polymerized before adding 90 kDa-Palld or Ig3-Palld; however, in our next set of experiments, palladin (90 kDa or Ig3) was incubated with G-actin under polymerizing or non-polymerizing conditions before low-speed centrifugation to separate crosslinked actin from filaments formed during polymerization in the presence of palladin. The data shown in Figure 3 reveals that both 90 kDa-Palld and Ig3-Palld are capable of polymerizing G-actin upon the addition of increasing concentration of palladin in both G- and F-buffer conditions. However, Ig3-Palld produces significantly more crosslinked or bundled actin filaments in the G-buffer condition (Fig. 3A) whereas 90 kDa-Palld generates the same amount of bundles under either buffer condition (Figure 3B). Ig3-Palld also increases the overall amount of polymerized actin as shown by the increased percentage of actin in the pellet fraction as compared to 90kDa-Palld.

**Figure 3.**
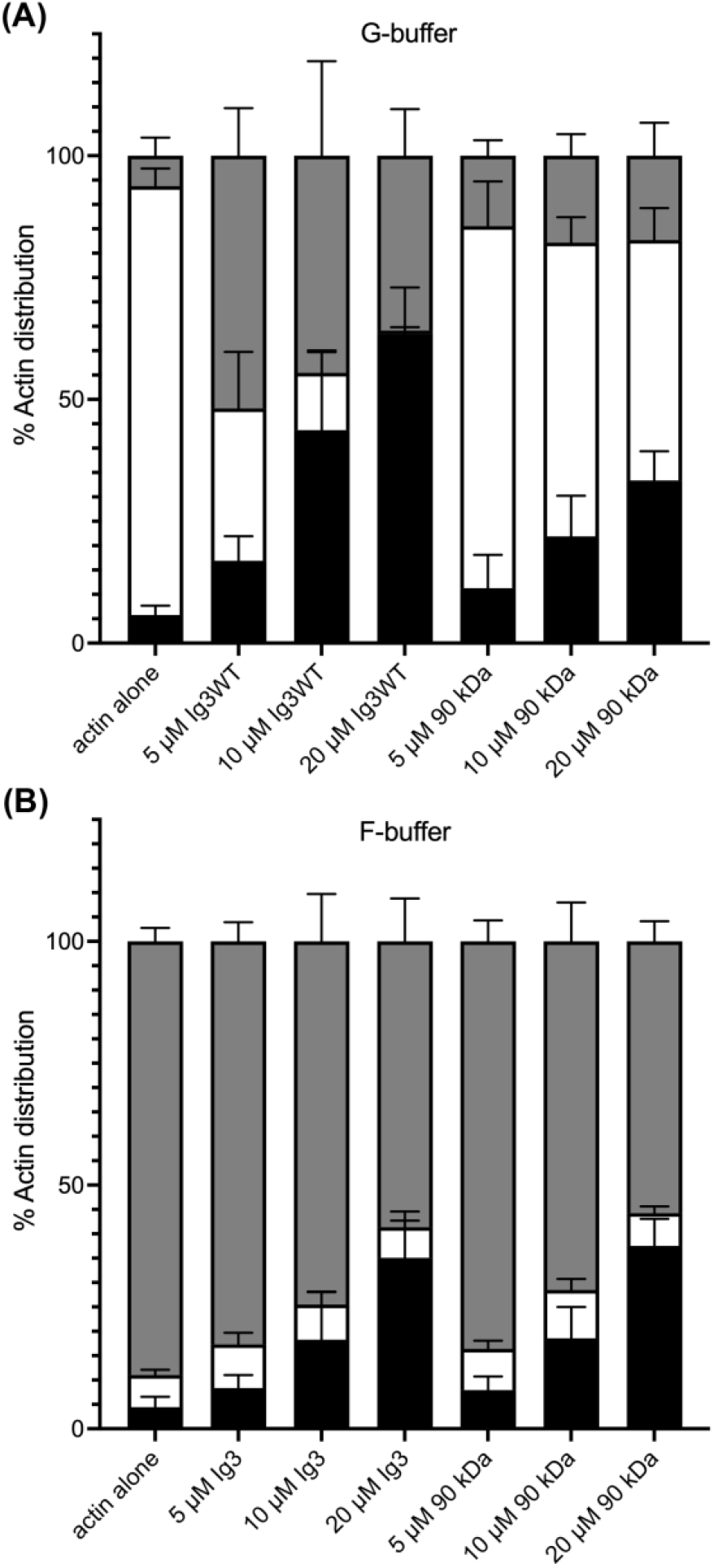
Co-polymerization bundling assay. G-actin was incubated with increasing amounts of either Ig3-Palld or 90 kDa-Palld under either G-Buffer (A) or F-Buffer (B) conditions. The percentage of actin found in the low-speed bundle is represented by the black bar, soluble portion is white, and high-speed pelleted actin is gray. Data are means +/- standard deviation for three or more separate measurements.

### 2.4 Actin polymerization by palladin

While the co-sedimentation assays helped us to quantify polymerization and bundling efficiency with both 90 kDa-Palld and Ig3-Palld under equilibrium conditions, we also wanted to directly measure the effect of 90 kDa-Palld and Ig3-Palld on the actin polymerization rate. The polymerization of pyrene-labeled G-actin (5%) was detected by monitoring the increase in fluorescence intensity which is proportional to the polymerization of G-actin into F-actin (Cooper et al. 1983). In this assay we measured the polymerization of G-actin (5 µM) in the presence and absence of various concentrations of either 90 kDa-Palld or Ig3-Palld (0-20 µM) under G-buffer conditions (Figure 4). The data shows that 90 kDa-Palld (10 µM) has a three-fold slower polymerization rate than Ig3-Palld (10 µM) (Table 2) and there is also an increase in the maximum intensity at the plateau for Ig3-Palld over that of 90 kDa-Palld. Keeping the previous co-polymerization assay in mind, the rational explanation of the increase of fluorescence intensity plateau is likely due to the Ig3 domain binding to G-actin which promotes actin polymerization while also forming actin bundles. These results are in agreement with the previous co-polymerization comparison assay where the 90 kDa-Palld displayed lower binding and bundling ability in the G-buffer condition than Ig3-Palld despite the fact that the F-actin binding affinity is greater for 90 kDa-Palld. One explanation for this data is that 90 kDa-Palld may autoinhibit and prevent the binding to *monomeric* actin which will be further examined in the discussion.

**Figure 4.**
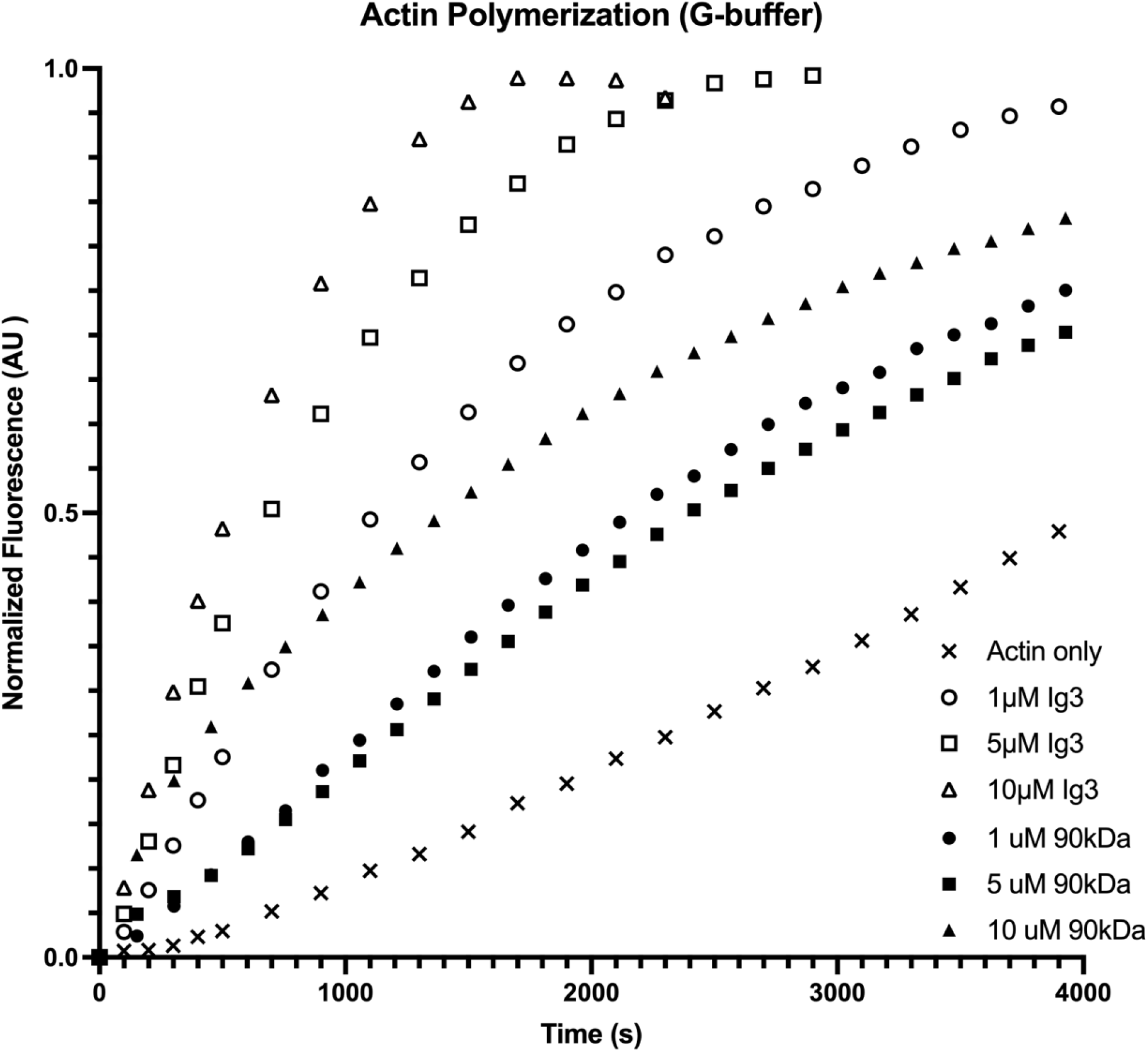
Fluorescence spectroscopy to monitor pyrene actin polymerization. Spontaneous assembly of 5% pyrene labeled G-actin in the presence of various concentration of either 90 kDa-Palld (closed symbols) or Ig3-Palld (open symbols) under G-buffer conditions. The fluorescence intensity was monitored to compare the polymerization rates of 90 kDa-Palld and Ig3-Palld.

**TABLE 2.** Polymerization rate of different concentration of the 90 kDa-Palld and Ig3-Palld

|  | 90 kDa-Palld Concentrations |  |  | Ig3-Palld Concentrations |  |  |  |
| --- | --- | --- | --- | --- | --- | --- | --- |
|  | 1 μM | 5 μM | 10 μM | 1 μM | 5 μM | 10 μM | 20 μM |
| Polymerization rate (nM/s) | 0.98 | 0.98 | 0.98 | 0.98 | 1.96 | 2.94 | 2.94 |

### 2.5 Palladin prevents actin depolymerization

To compare the effect of 90 kDa-Palld with Ig3-Palld on actin filament stability, we measured the disassembly rate of pyrene-labeled actin filaments in the presence and absence of various concentrations of 90 kDa-Palld or Ig3-Palld (0-20 μM) and Latrunculin A (10 μM LatA), an actin monomer sequestering agent (Coue et al. 1987; Yarmola et al. 2000). As expected, the fluorescence intensity decreased in a time-dependent manner in the presence of LatA alone indicating net actin filament depolymerization (Figure 5). We calculated the disassembly rate of the actin filaments from the initial slope of the curves to reveal that the presence of LatA increases the disassembly rate by 12-fold when compared to actin filaments alone (Table 2). Addition of Ig3-Palld significantly decreased the disassembly rate in a dose-dependent pattern. For the highest concentration of Ig3-Palld (20 μM), the rate of LatA-induced disassembly was decreased by 12-fold compared to actin alone. However, 90 kDa-Palld also decreased the depolymerization rate by 12-fold compared to actin alone at all concentrations tested. These results indicate the ability of 90 kDa-Palld to bind tightly to actin filaments and prevent the LatA depolymerization effect. We compared the effect of 90 kDa-Palld with that of Ig3-Palld in the stabilization of actin filaments and found that at high concentrations (10 and 20 μM) both 90 kDa-Palld and Ig3-Palld promote stability of the actin filaments to a similar degree. However, 90 kDa-Palld stabilizes the actin filament almost four times more than the Ig3 domain at low protein concentrations (2 and 5 μM). Low concentrations of 90 kDa-Palld protect the filaments from depolymerization and decrease the dissociation of monomers from filaments to a greater degree than Ig3-Palld.

**Figure 5.**
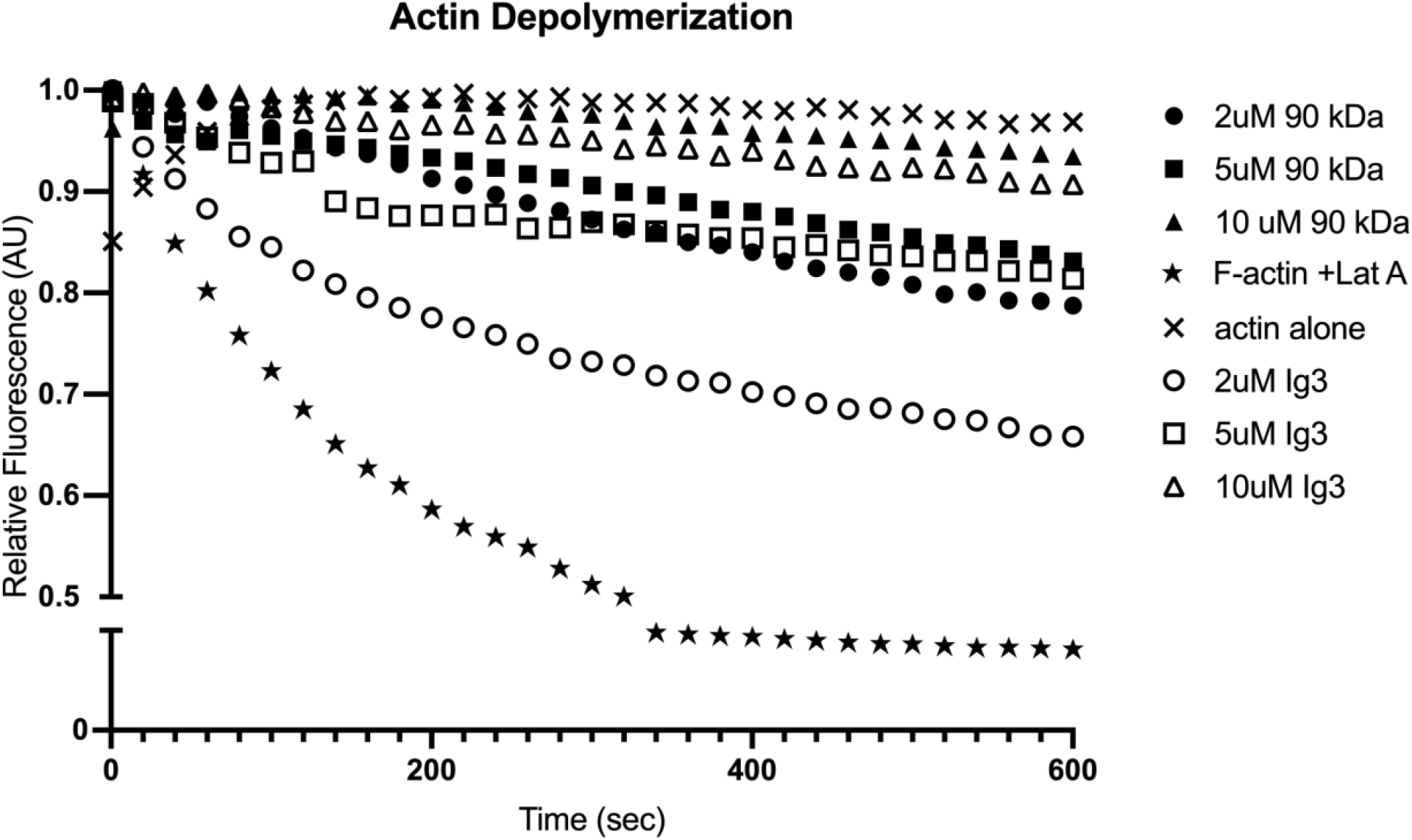
90 kDa-Palld and Ig3-Palld stabilize actin filaments. G-actin (2 μM, 5% pyrene labeled) was preassembled to filaments in polymerization buffer for one hour. F-actin was then incubated with various concentrations of 90 kDa-Palld (closed symbols) or Ig3-Palld (open symbols) (2-10 μM) for half an hour. Latrunculin A was then added to each reaction mixture at time point 0, and fluorescence measurement began immediately. Fluorescence of F-actin alone (x) and F-actin with Lat A only (stars) were also measured as controls. The signal of the fluorescence was normalized so that the fluorescence of G-actin was 0 and F-actin was 100.

### 2.6 Monomeric actin binding by palladin

Supported by the actin polymerization and copolymerization data presented in this study as well as previous work (Gurung et al. 2016), we posited that Ig3-Palld would bind to monomeric actin in order to promote nucleation. In this monomeric actin binding assay, three columns of AffiGel 10 resin were utilized for each protein—a control column with no actin bound, a second column with G-actin bound, and a third column with F-actin bound (Miller et al. 1989; Nefsky et al. 1989). Assays were performed in triplicate by adding purified palladin [Ig3WT, tandem Ig34WT, Ig3(K15A, K18A, K51A), or 90 kDa], rinsing unbound protein with wash buffer, and finally eluting any bound protein. All control columns showed little bound palladin in the elution fractions, indicating that palladin does not bind nonspecifically to the resin. As expected, both wildtype Ig3 and Ig34 show significant binding to the F-actin column as shown in Figure 6. Ig34 is known to bind to F-actin more tightly than the isolated Ig3 domain which is also reflected by the increased binding seen in these results (Dixon et al. 2008). Both of these proteins also show equal or greater interaction with the G-actin column, supporting the possible role of palladin as an actin nucleator. However, 90 kDa-Palld shows a much lower degree of bound palladin, comparable to that of the actin binding mutant of Ig3 (K15A, K18A, K51A) which has been shown to have diminished actin binding ability in previous co-sedimentation assays (Beck et al. 2013). Since 90 kDa-Palld is already known to bind to F-actin, the low levels of interaction indicated by this assay support the concept 90 kDa-Palld may be autoinhibited under these conditions.

**Figure 6.**
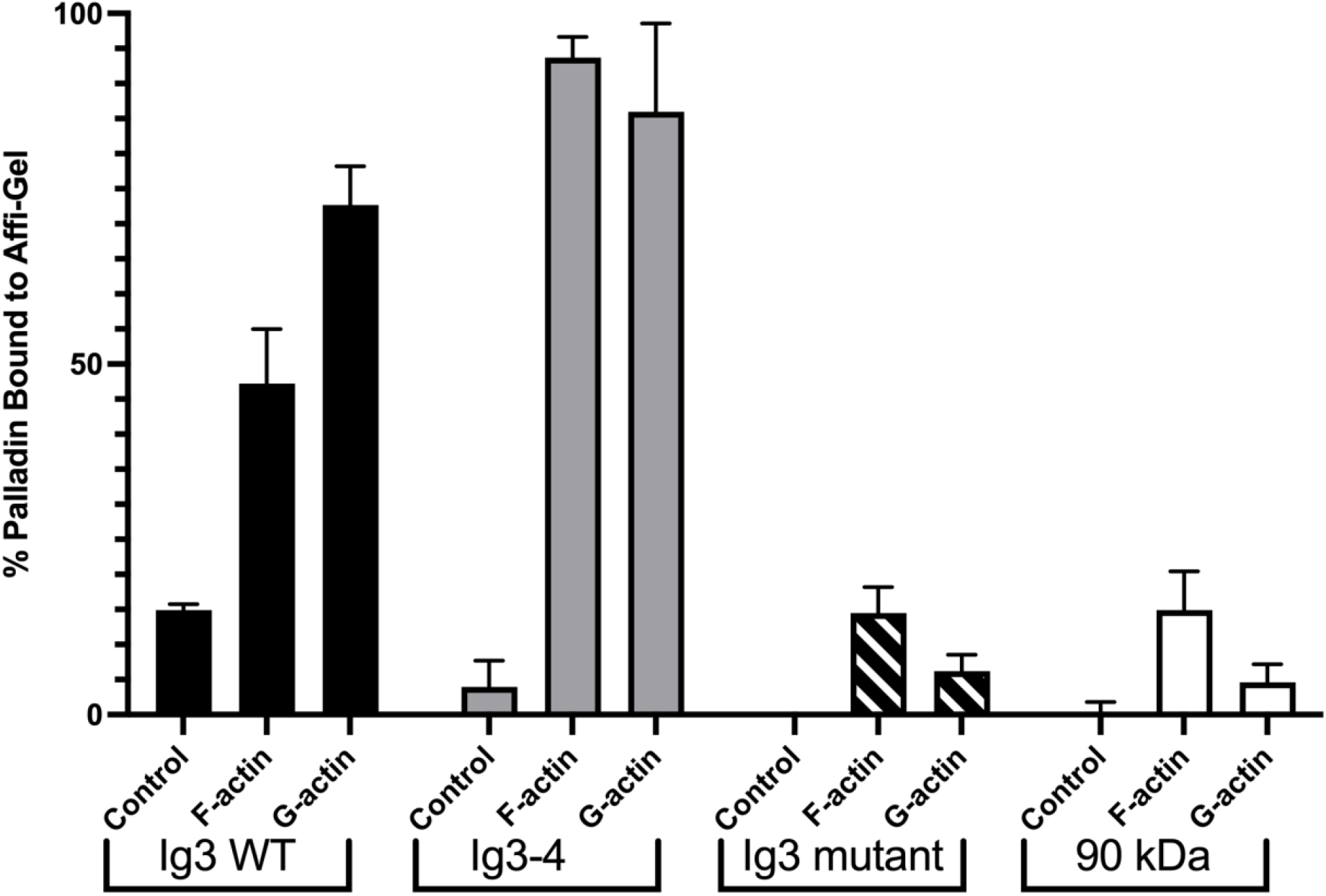
Affinity column comparison of G-versus F-actin binding. Percent of palladin bound to Affi-Gel 10 under three separate conditions. Columns were prepared as Control, F-actin, or G-actin containing no, filamentous, or monomeric actin, respectively. For each palladin protein, bars represent the percent of protein found in the elution or bound fraction. Wildtype Ig3-Palld (black) and Ig3-4-Palld tandem domain (grey) indicate binding to both F- and G-actin. The Ig3(K15,18,51A)-Palld mutant shows diminished actin binding ability (striped) and 90kDa-Palld (white) shows minimal binding to any column. Error bars indicate standard deviation for three or more separate measurements.

## 3 DISCUSSION

### 3.1 Comparison of F-actin binding and bundling

The aim of this study was to determine the effect of 90 kDa-Palld, which is expressed in all tissues, on the assembly and disassembly of actin filaments and to compare the results of the 90 kDa isoform 4 with the isolated actin binding domain of palladin (Ig3-Palld). We also sought to compare the binding affinity of 90 kDa-Palld purified from bacteria cells with that previously measured for 90 kDa-Palld purified from insect cells (Dixon et al. 2008). We found that our binding affinity data (K_d_ = 3.1 ± 0.9 μM) was in close agreement with the previously measured data from the insect cells (K_d_ = 2.1 ± 0.5 μM) (Dixon et al. 2008). By determining the dissociation constant from the binding curves, we found that 90 kDa-Palld has a slightly tighter binding affinity than Ig3-Palld with 2.11 ± 1.09 μM and 3.42 ± 1.24 μM values, respectively. A previous study had shown that the Ig34 tandem domains of palladin enhance the binding affinity for F-actin compared to the Ig3 domain alone (Dixon et al. 2008). This indicates that there are other regions of 90 kDa palladin that contribute to the interactions with F-actin which explains the higher binding affinity of 90 kDa-Palld compared to the isolated Ig3-Palld domain. In addition to the F-actin binding affinity, we also compared the bundling ability of 90 kDa-Palld with our previously measured results for Ig3-Palld (Gurung et al. 2016). In the bundling assay, we found that 90 kDa-Palld has a significantly higher bundling capacity than Ig3-Palld. For example, at 10 μM 90 kDa-Palld almost 50 % of actin filaments were bundled while at 10 μM Ig3-Palld only ∼20 % of actin filaments were bundled. Furthermore, the data shows that bundling by the 90 kDa-Palld is saturated at a 1:1 ratio, which is not the case with Ig3-Palld.

### 3.2 Comparison of G-actin binding

From previous co-polymerization studies in the Beck lab, we found that palladin is able to facilitate actin polymerization under non-polymerizing conditions in the absence of ionic strength that is known to enhance actin polymerization (Gurung et al. 2016). Here, this same co-polymerization assay was carried out to compare the ability of 90 kDa-Palld with that of the isolated Ig3-Palld domain to promote the polymerization of G-actin under both F-buffer and G-buffer conditions. We found that 90 kDa-Palld generates the same amount of bundling of actin filaments under polymerization conditions (F-buffer) as compared to Ig3-Palld. However, Ig3-Palld has a larger effect in bundling the monomeric actin into crosslinked filaments under non-polymerization conditions (G-buffer) than 90 kDa-Palld and produces more polymerized actin as seen by the increased pellet fraction. This data indicates that Ig3-Palld may have a higher binding affinity for the monomeric actin than 90 kDa-Palld which is also further supported by the results of the actin monomer binding assay with the AffiGel 10 resin.

In the affinity binding assay, both Ig3-Palld and the tandem Ig34-Palld showed high binding levels to F-actin compared to the control columns as expected. Both of these proteins also showed binding to G-actin monomers. Several lysine mutations in Ig3 (K15A, K18A, K51A) have previously been shown to interfere with F-actin binding ability (Beck et al. 2013). This G-actin binding assay also showed decreased interaction with this mutant as compared to the wildtype Ig3 domain. Furthermore, the lack of G-actin binding also indicates that these three lysine residues may be involved with binding to monomeric actin. Similarly, 90 kDa-Palld shows low levels of binding to both F- and G-actin. Autoinhibition could be an explanation for these results if the binding domain in 90 kDa-Palld is hindered from interacting with the actin in these conditions.

### 3.3 Comparison of polymerization and depolymerization

There were no previous investigations on how 90 kDa-Palld affects the rate of actin polymerization or the effects on stabilizing actin filaments. Here, we directly measured the effect of 90 kDa-Palld on the polymerization rate of G-actin and compared the rate with Ig3-Palld (Gurung et al. 2016). Our kinetics data shows that 90 kDa-Palld also regulates the polymerization dynamic of actin filaments directly. Surprisingly, Ig3-Palld (10 µM) has a four-fold greater polymerization rate than 90 kDa-Palld (10 µM). In addition, we found that Ig3-Palld significantly promotes actin assembly with increasing protein concentrations. However, increasing concentrations of 90 kDa-Palld does not change the effect on the polymerization rate of actin which may indicate that the actin polymerization by 90 kDa-Palld is already saturated at the concentrations used in these assays (1, 5 and 10 µM). The co-polymerization assay indicates that both 90 kDa-Palld and Ig3-Palld produced similar amounts of actin filaments under F-buffer conditions but that only Ig3 enhances bundling of actin under G-actin buffer conditions. In addition, the polymerization rate of the 90 kDa-Palld is 4-fold slower than Ig3-Palld which suggests that 90 kDa-Palld has tighter affinity for F-actin than G-actin. These findings suggest that 90 kDa-Palld may be autoinhibited in such a manner that blocks the binding site for monomeric actin while maintaining F-actin binding capacity.

To determine whether 90 kDa-Palld contributes to the stability of actin filaments, we conducted a series of LatA-induced depolymerization assays. In these assays, we directly compared the effect of various concentrations of 90 kDa-Palld and Ig3-Palld on the depolymerization rate of actin filaments. The data shows that the low concentration (2 and 5 µM) of 90 kDa-Palld has a four-fold greater effect on actin filament stability compared to Ig3-Palld. However, higher concentrations (10 and 20 µM) of both 90 kDa-Palld and Ig3-Palld equally stabilize the actin filaments.

### 3.4 Model of autoinhibition in 90 kDa palladin

Our data indicates that 90 kDa-Palld binds tightly and crosslinks F-actin but does not interact significantly with G-actin as opposed to the isolated Ig3-Palld domain. Based on our co-polymerization and polymerization assays, these findings support the hypothesis that 90 kDa-Palld may be autoinhibited thus blocking the G-actin binding site and preventing the binding of 90 kDa-Palld with monomeric actin. Our proposed models in Figure 7 indicate that the C-terminus and N-terminus of 90 kDa-Palld may interact with each other to form the closed conformation. In both models, this closed conformation would therefore block the Ig3-Palld domain from binding with monomeric actin. Another possibility is that some currently unknown cause for a conformational change of 90 kDa-Palld triggers the closed form into an open conformation that can then bind to the F-actin and that binding to G-actin may not induce this conformational change. In other words, 90 kDa-Palld stays in its closed conformation which prevents it from binding to monomeric actin.

**Figure 7.**
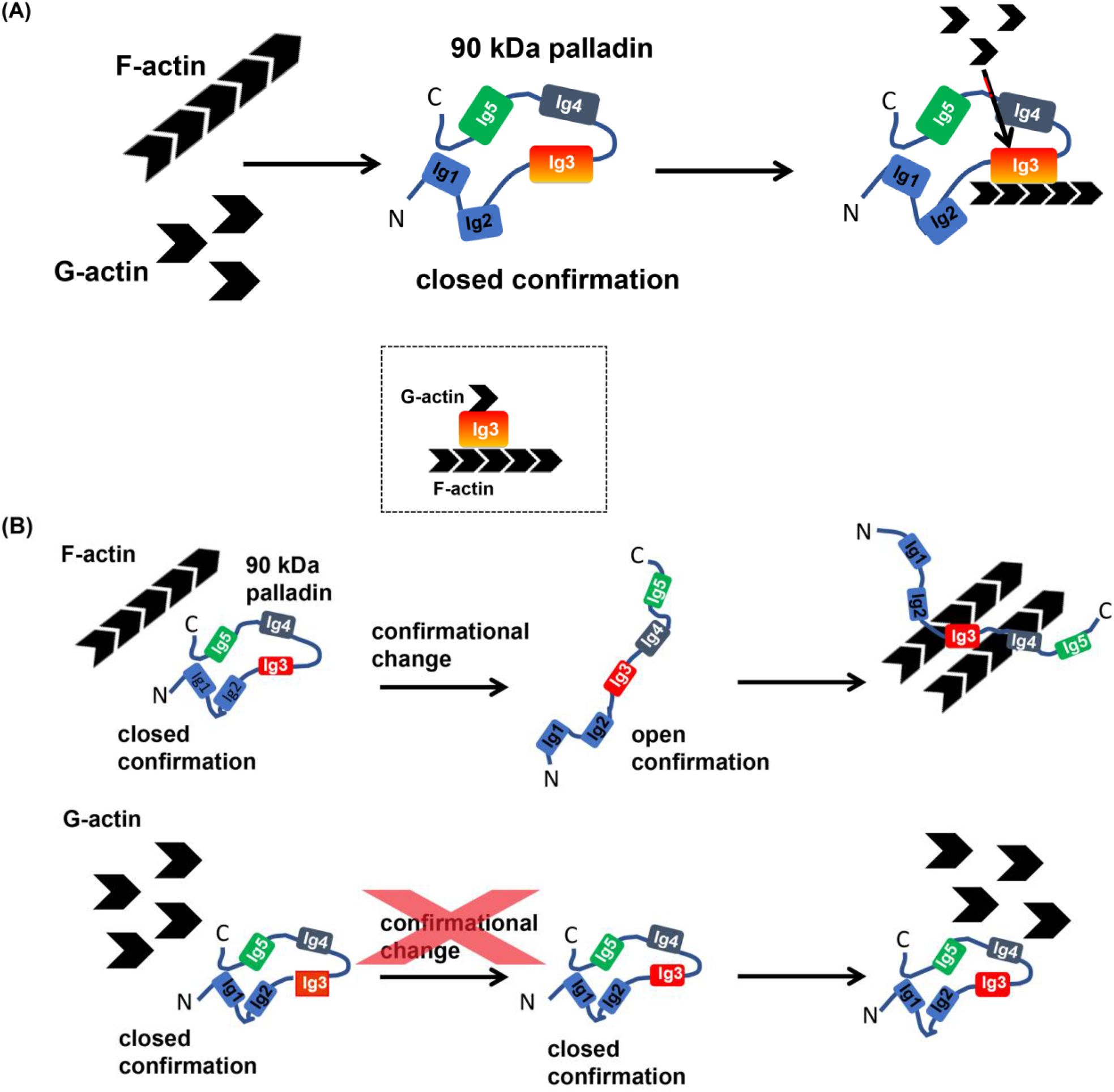
Proposed models for the autoinhibition of 90 kDa-Palld. (A) Interactions between the C- and N-terminus of 90 kDa-Palld forms a putative closed conformation which prevents the binding of the Ig3 domain to G-actin. (B) Alternatively, 90 kDa-Palld may exist in a closed form that can undergo conformational changes upon binding F-actin or some other regulatory protein (top row), whereas no conformational change occurs in the presence of the G-actin (bottom row).

An autoinhibition function has been previously described for the actin binding protein vinculin. Vinculin is a globular protein that has five helical domains: four head domains (D1-D4) and the vinculin tail domain (Vt) (Ziegler et al. 2006). The Vt domain is known to be the actin binding site, similar to Ig3 in palladin. Vt also binds to the vinculin head domain to form the closed conformation of vinculin (Carisey et al. 2011). This closed conformation of vinculin autoinhibits vinculin from binding to F-actin (Nhieu et al. 2007). Further investigations need to be carried out to establish this proposed autoinhibition model for palladin.

## 4 CONCLUSIONS

Previous research has established that the Ig3-Palld domain is the minimal binding site for F-actin (Dixon et al. 2008). Recently, Ig3-Palld has been shown to increase the polymerization rate and decrease the disassembly rate of actin filaments (Gurung et al. 2016). While previous research has been carried out with 90 kDa-Palld that was purified from insect cells, this was limited to F-actin binding and bundling assays only (Dixon et al. 2008). In our previous work, we established a role for palladin in the generation of actin-rich structures during *L. monocytogenes* infections (Dhanda et al. 2018). We found that palladin is crucial for the structural maintenance of *L. monocytogenes* actin-rich comet tails and revealed that there are specific regions of palladin, such as the actin binding and the VASP binding sites, that are required to maintain proper morphology of the *L. monocytogenes* actin comet tails. The most significant discovery in this previous study was that only the 90 kDa-Palld, but not the Ig3-Palld domain, can compensate for Arp2/3 complex during *L. monocytogenes* motility indicating that 90 kDa-Palld can form the necessary branched or meshwork structures needed for highly motile cells.

In this study, we have been able to successfully purify 90 kDa-Palld, or isoform 4, from bacterial cells for the first time. This isoform is ubiquitously expressed in all cell types and is specifically upregulated in cancer cells (Welsch et al. 2007; Goicoechea et al. 2009). Here we compared the effect of 90 kDa-Palld and Ig3-Palld on actin binding, bundling, and dynamics. This work has demonstrated that 90 kDa-Palld has a higher binding affinity and bundling capacity for actin filaments than isolated Ig3-Palld. In addition, we have shown that 90 kDa-Palld enhances the actin polymerization rate but to a lesser degree than Ig3-Palld. This suggests that 90kDa-Palld may be autoinhibited and is thereby blocking the binding site for monomeric actin. In future research, autoinhibition of 90 kDa palladin could be verified by measuring the binding affinity between different purified fragments of the protein.

## 5 MATERIALS AND METHODS

### 5.1 Protein Expression and Purification

The Ig3 domain of palladin (Ig3-Palld) was purified as previously described (Gurung et al. 2016). The 90 kDa isoform of palladin (90kDa-Palld) in the phrGFP II N (4.9 kb) mammalian expression vector (gift from C.A. Otey) was subcloned into the pTBSG bacterial expression vector to express the protein in *E. coli*. The pTBSG-90kDa-Palld was transformed into BL21 (DE3)-RIPL *E. coli* cells (Agilent Technologies) (Qin et al. 2008). The 90 kDa-Palld was purified as previously described for Ig3-Palld (Dixon et al. 2008) after overexpression in autoinduction media (Studier 2005) with a few differences. Expression was carried out at 15 ºC, as opposed to 18 ºC, with shaking for ∼15 hours because the slightly lower temperature increased the yield of soluble protein. After harvesting the cells, the pellet was resuspended in 50 ml of lysis buffer (50 mM sodium phosphate pH 7.4, 300 mM NaCl, 10% glycerol, 3 mM DTT) for 8 L of growth and protease inhibitor cocktail (Cat. No. P8849, Sigma-Aldrich) was added. Then, a PROTEINDEX™ Ni-Penta™ Agarose 6 Fast Flow column (Marvelgent Biosciences, Inc.) was used to purify the supernatant fraction after lysis and centrifugation. The supernatant was incubated overnight with rotation at 4 ºC with the nickel-column pre-equilibrated into lysis buffer. Next, the column was washed with 3-5 column volumes of wash buffer (50 mM sodium phosphate pH 7.4, 300 mM NaCl, 10% glycerol, 3 mM DTT, 10 mM imidazole). The protein was eluted using elution buffer (50 mM sodium phosphate pH 7.4, 300 mM NaCl, 10% glycerol, 3 mM DTT, 300 mM imidazole). All PROTEINDEX™ Ni-Penta™ Agarose 6 Fast Flow column purification was performed at 4 ºC to prevent protein degradation. A 10% SDS-PAGE gel was used to confirm the presence of the protein in the elution fractions. Then, the protein was dialyzed into 1 L lysis buffer overnight at 4 ºC to remove the high imidazole concentration. The protein was then dialyzed in 4 L of S-column buffer A (25 mM KH_2_PO_4_ pH 6.5, 50 mM NaCl, 2 mM DTT) until the protein pH had been lowered to 6.5 for no more than 6 hours. Then, this protein fraction was further purified using the cation exchange column as previously described in the Ig3 purification (Dixon et al. 2008). The peak fractions were collected and the presence of 90 kDa-Palld was confirmed by running in 10% SDS-PAGE gel. After purification, 90 kDa-Palld was dialyzed into the storage buffer (20 mM HEPES, pH 7.5, 150 mM NaCl, 5 mM DTT) and stored at 4 °C to be used within one week. To measure the concentration of 90 kDa-Palld the extinction coefficient of 55,975 M^−1^cm^−1^ was used at 280 nm. Reagents were purchased from Fisher Scientific unless otherwise noted.

### 5.2 Actin preparation from rabbit muscle and labeling

Actin was purified from rabbit muscle acetone powder (Pel-Freez Biologicals) as previously described (Spudich et al. 1971). N’-[3-pyrenyl]-maleimide (M.W. 297.3, Sigma-Aldrich) was used to react with the gel-filtered G-actin as previously described (Cooper et al. 1983).

### 5.3 Actin co-sedimentation assays

Three different actin co-sedimentation assays were utilized to monitor bulk interactions between actin and palladin. Two of these assays employed F-actin to measure binding or crosslinking efficiency of 90 kDa-Palld and Ig3-Palld (Dixon et al. 2008). In these assays, actin was pre-polymerized with the polymerization buffer (10 mM Tris, pH 8.0, 100 mM KCl, 2 mM MgCl_2_ and 2 mM DTT) and then 10 µM palladin (90 kDa or Ig3) was incubated with varying concentrations of F-actin (0-25 µM). To monitor F-actin binding, samples were then centrifuged at 100,000 *xg* for 30 minutes to sediment F-actin and any interacting proteins. For bundling, or crosslinking, assays the samples were first centrifuged at 5,000 *xg* for 10 minutes followed by separation of the supernatant from the pellet (bundle). The supernatant samples were then centrifuged at 100,000 *xg* for 30 minutes to sediment remaining actin filaments in the pellet from proteins in the supernatant. The co-polymerization assays were carried out to observe whether incubating the 90 kDa-Palld or Ig3-Palld with G-actin during polymerization affects the amount of crosslinked actin formed. Here, G-actin (10 μM) was incubated with various concentrations of Ig3-Palld or 90kDa-Palld (0-20 μM) under either polymerization conditions of F-buffer (5 mM Tris-HCl, pH 8.0, 100 mM KCl, 2 mM MgCl_2_ supplemented with 1 mM DTT and 0.2 mM ATP) or under non-polymerization conditions of G-buffer (5 mM Tris-HCl, pH 8, 0.1 mM CaCl_2_, 0.2 mM DTT, 0.2 mM ATP, 0.02% NaN_3_). Palladin was in either G-buffer for Ig3-Palld or in storage buffer for 90 kDa-Palld, where both buffers do not contain the MgCl_2_ or KCl necessary for initiation of actin polymerization. This mixture was allowed to co-incubate for one hour and then centrifuged at 5,000 x*g* for 10 min to pellet the bundled F-actin. To pellet the actin filaments, the supernatant was then centrifuged at 100,000 x*g* for 30 minutes. In all three assays, the fractions (supernatant, pellet, and bundle) were separated by 10% or 15% SDS-PAGE. Gels were scanned and ImageJ software (Abràmoff et al. 2004) was used to quantify the percent of actin in the fractions for bundling or co-polymerization assays or percentage of fractionated palladin in binding assays.

### 5.4 Pyrene fluorescence assay of actin polymerization

N’-[3-pyrenyl]-maleimide (Sigma-Aldrich) was used to react with the gel-filtered G-actin as previously described (Cooper et al. 1983). On the day of the experiment, a 10X priming solution (10 mM EGTA and 1 mM MgCl_2_) was added to the 5% pyrene/actin stock solution to convert Ca-G-actin into Mg-G-actin for 2 minutes. Then, the stock solution was incubated with various concentrations of palladin proteins (Palld-Ig3 or 90 kDa-Palld) in G-buffer conditions (without KCl). Polymerization of G-actin was quantified with excitation at 365 nm and emission at 385 nm on a fluorescence spectrophotometer (PTI). To ensure that the palladin storage buffer had no effect on the polymerization rate, a control sample was utilized wherein the same volume of palladin storage buffer was used instead of the protein. For data quantification, raw data were normalized by subtracting the baseline fluorescence and then dividing the fluorescence values by the steady-state plateau fluorescence. By assuming that at equilibrium the total concentration of polymers is equal to the total concentration of actin minus the critical concentration, we can measure the overall polymerization rate of each polymerization curve under G-buffer condition by plotting the slope of the linear region of each concentration curve and converting the relative fluorescence units/s into nM actin/s.

### 5.5 Actin depolymerization

The stock solution of 10 μM of 5% pyrene-labeled G-actin was incubated at room temperature with MKEI polymerization buffer plus 1 mM DTT and 0.2 mM ATP for one hour to make 10 μM of 5% pyrene-labeled F-actin. In 200 μl reactions, 2 μM 5% pyrene-labeled F-actin was incubated with various concentrations of Ig3-Palld or 90 kDa-Palld (1-20 μM) at room temperature for 30 minutes. After incubation, 2 μl of 1 mM Latrunculin A (Calbiochem) was added to the reaction mixture and actin filaments disassembly was immediately measured using pyrene fluorescence for 600 s. Then, the fluorescence emission at 385 nm (excitation at 365 nm) of each sample was normalized to 1 AU at steady state by dividing the first point of fluorescence intensity to the rest of the polymerization curve.

### 5.6 Affinity Chromatography

This procedure was adapted from two previously published protocols (Miller et al. 1989; Nefsky et al. 1989). Monomeric actin was dialyzed to G-Buffer (5 mM HEPES, pH 7.5, 0.2 mM CaCl_2_, 0.2 mM ATP) and then centrifuged at 100,000 *xg* for 2 hours to remove polymers. The supernatant was used directly for monomer binding chromatography. For polymer binding chromatography, actin was polymerized by addition of an equal volume of 2x F-Buffer (100 mM HEPES, pH 7.5, 0.2 M KCl, 4 mM MgCl_2_). For F-actin columns, polymerized actin was added to the Affi-Gel 10 resin (BioRad) and allowed to couple for 1-15 hours at 4 °C. The reaction was terminated by addition of 3 M monoethanolamine, pH 8.0 at a final concentration of 50 mM to block active esters. The column was washed with F-Buffer to remove unbound actin. The column was washed with 3-5 CV of Wash Buffer (50 mM HEPES, pH 7.5, 1 M KCl, 2 mM MgCl_2_). F-actin columns were stored in F-Buffer containing 10 mg of phalloidin per mL and 0.02% NaN_3_ at 4 °C and reused for up to 3 weeks. For G-actin columns, the Affi-Gel 10 resin was partially inactivated by incubation for 1.5 hours in G-Buffer to prevent denaturation of G-actin. Monomeric actin was added to the column and allowed to couple for 20 minutes before the addition of 50 mM monoethanolamine. The column was washed with G-Buffer supplemented with 1 M NaCl followed by G-Buffer supplemented with 5 mM ATP. G-actin columns were prepared on day of use. A control column without actin was used to monitor palladin binding to the resin. Control columns were prepared as G-actin columns without the actin addition and coupling step. All columns were first equilibrated to Buffer A (2 mM Tris-HCl, pH 8.0, 2 mM CaCl_2_, MgCl_2_, 0.1 mM NaN_3_, 10% glycerol) before being loaded with the actin binding protein and allowed to couple for 30 minutes. The flow through was collected and the column was washed with A-Buffer to remove any unbound actin binding protein. Elution of the actin binding protein was performed with A-Buffer supplemented with (0.1-1.0 M KCl or 1mM ATP + 3mM MgCl_2_). Each assay was performed in triplicate and the overall fraction of bound actin binding protein was determined by ImageJ analysis (Abràmoff et al. 2004).

## Acknowledgements

This research was financially supported by NIH AREA Award R15GM120670 to MRB. We thank Beck Lab undergraduates Hannah Rai and Kelsey Young for assisting with several experiments.

## Author Contributions

SA, RK, and OA performed protein expression and purification, SA and OA collected actin binding and bundling data, SA also collected and analyzed fluorescence data, RK collected and analyzed monomer binding data, all authors contributed to the writing of the manuscript; SA, RK, OA, and MRB prepared the figures. MRB conceived and designed the experiments employed, supervised the research, secured funding and helped analyze the data.

