## Supplementary Figure S1 for "Functional comparison of full-length palladin to isolated actin binding domain"

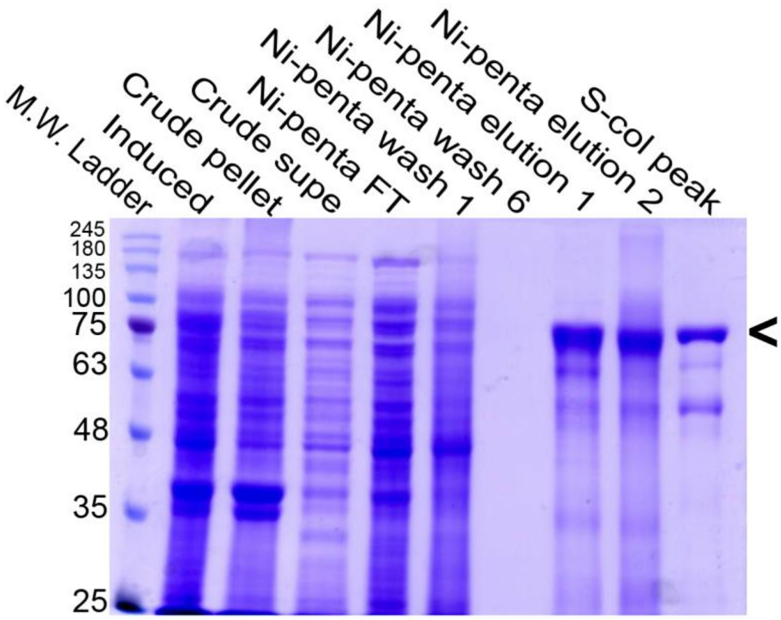


**Supplemental Figure S1.** SDS-PAGE gel showing steps of 90 kDa-Palld purification from *E. coli*. Ten percent SDS-PAGE gel stained with Coomassie blue. Lane 1 contains BLUEstain 2 protein ladder (Gold Biotechnology). Lane 2 is whole cell lysate of bacteria expressing 90 kDa palladin. Lanes 3 and 4 contains the insoluble and soluble protein fractions after sonication and centrifugation. Lanes 5-9 show the His-tagged protein fractions from Ni-Penta™ (Marvelgent) affinity column. Lane 10 contains the pure 90 kDa-Palld after purification over the cation exchange column. Carrot to right indicates size of band for 90 kDa-Pallad.
